# The joint evolution of lifespan and self-fertilisation

**DOI:** 10.1101/420877

**Authors:** Thomas Lesaffre, Sylvain Billiard

## Abstract

In Angiosperms, there exists a strong association between mating system and lifespan. Most self-fertilising species are short-lived and most predominant or obligate outcrossers are long-lived. This association is generally explained by the influence of lifespan on the evolution of the mating system, considering lifespan as fixed. Yet, lifespan can itself evolve, and the mating system may as well influence the evolution of lifespan, as is suggested by joint evolutionary shifts of lifespan and mating system between sister species. In this paper, we build modifier models to study the joint evolution of self-fertilisation and lifespan, including both juvenile and adult inbreeding depression. We show that self-fertilisation is expected to promote evolution towards shorter lifespan, and that the range of conditions under which selfing can evolve rapidly shrinks as lifespan increases. We study the effects of inbreeding depression affecting various steps in the life cycle, and discuss how extrinsic mortality conditions are expected to affect evolutionary associations. In particular, we show that selfers may sometimes remain short-lived even in a very stable habitat, as a strategy to avoid the deleterious effects of inbreeding.

## 1 Introduction

In Angiosperms, strong associations exist between mating systems and other life-history traits, such as dispersal [3], allocation to male *vs.* female functions [7], or lifespan [4]. The latter association has received limited attention. Indeed, while it has long been recognised that most self-fertilising species are short-lived and most predominant or obligate outcrossers are long-lived [4, 12, 25, 32], relatively few arguments have been advanced to explain this association. Stebbins [32] proposed that the reproductive assurance provided by self-fertilisation could be less beneficial to perennials than it is to annuals, since they get more than one try at reproducing. Later, Lloyd [21] suggested that self-fertilisation in perennials could cause the consumption of resources that could have been more advantageously allocated to post-breeding survival or future outcrossed reproduction (between-seasons seed discounting). Morgan, Schoen, and Bataillon [24] investigated the validity of these arguments by developing a phenotypic model and concluded that the association between annuality and selfing is more likely to be due to between-seasons seed discounting, rather than reproductive assurance. They also showed the importance of the repeated effect of adult inbreeding depression for the maintenance of outcrossing in perennials. Empirical evidence also suggests that inbreeding depression is overall higher in perennials than in annuals [2], which could constitute an additionnal barrier to the evolution of self-fertilisation in perennials.

These arguments focus on the consequences of perenniality for the evolution of self-fertilisation, considering lifespan as a fixed characteristic. Yet, lifespans evolve in nature [31], and joint evolutionary shifts of lifespan and mating system have been documented. Indeed, the transition to self-fertilisation is often associated with significant morphological changes, such as vegetative size and flower size reduction [the selfing syndrome, 29], and lifespan shortening compared to outcrossing relatives [14]. For example, *Arabidopsis thaliana* is a highly selfing annual that recently differentiated from its self-incompatible and perennial relatives *A. lyrata* and *A. halleri* [11]. Furthermore, studying pairs of sister species, Grossenbacher, Briscoe Runquist, Goldberg, and Brandvain [16] found numerous joint shifts towards selfing and annuality from outcrossing, perennial ancestors in genus such as *Mimulus* or *Medicago*, and very few shifts to longer lifespans in association with selfing. In fact, the only such shifts they found were observed in the *Oenothera* genus, where segregation and recombination are suppressed when reproducing by self-fertilisation, which implies that selfing individuals are effectively reproducing clonally [18].

These examples show that joint shifts of mating system and lifespan almost always occur in the same direction, that is towards lifespan shortening in association with selfing from outcrossing ancestors with longer lifespans. In such situations, lifespan shortening could have allowed for the evolution of self-fertilisation. Alternatively, the transition to self-fertilisation could have induced evolution towards a shorter lifespan. This possibility has seldom been investigated. Indeed, classical studies of the evolution of lifespan predict that it should be fine tuned to best fit the extrinsic mortality conditions experienced by the considered population, through optimal allocation of resources to reproduction, growth or survival [e.g. 10, 27] and senescence [e.g. 30], but rarely consider the evolution of lifespan in interaction with other traits. From a theoretical standpoint, the only study that, to our knowledge, investigated the influence of the mating system on lifespan evolution is that of Zhang [33], who developed a phenotypic model for the joint evolution of reproductive effort and sex allocation in partially self-fertilising hermaphrodites. Assuming a survival *vs.* reproduction trade-off [31], they reached the conclusion that reproductive effort increases (and lifespan decreases) when the selfing rate increases through greater allocation to the female function, provided that inbreeding depression is weak 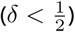, female reproduction is very costly, and juvenile survivorship is constrained within a narrow range of values. Importantly however, Zhang [33] assumed inbreeding depression to only affect the survival of juveniles to maturity, although inbreeding depression typically occurs over all stages of the life cycle [17]. In summary, on the one hand, the influence of lifespan on the evolution of the mating system has been studied, considering lifespan as a fixed characteristic [24]. On the other hand, the potential influence of the mating system on the evolution of lifespan has only been scarcely investigated, assuming no inbreeding depression occured in adults [33]. Finally, the question of the joint evolution of lifespan and mating system has never been tackled.

In this paper, we build modifier models [19] to investigate the joint evolution of lifespan and selfing, including inbreeding depression affecting various steps in individuals’ life cycle as fixed parameters. Following previous authors, we model the evolution of lifespan through that of reproductive effort, assuming a survival *vs* reproduction trade-off for which we consider various shapes [31], and incorporate extrinsic mortality as a constant parameter [27, 33]. We first study the evolution of each trait separately, taking the other as fixed, and incorporate inbreeding depression affecting juvenile and adult survival. In each case, we obtain accurate analytical approximations for the evolutionary stable strategies. We show that self-fertilisation is expected to favour evolution towards shorter lifespans when inbreeding depression affects adult survival. Conversely, we show that the range of inbreeding depression under which selfing can evolve rapidly shrinks as lifespan increases, in agreement with previous work [24]. Then, using individual-centered simulations along with our previous analytical approximations, we study the joint evolution of lifespan and selfing. We study the effects of inbreeding depression affecting various steps in the life cycle, and discuss how extrinsic mortality conditions are expected to affect evolutionary associations. In particular, we show that selfers may sometimes remain short-lived even in a very stable habitat, as a strategy to avoid the deleterious effects of inbreeding.

## 2 Methods

### 2.1 Outline of the model

#### Life cycle and demographic assumptions

We consider a very large population made of partially selfing hermaphrodites, which are assumed to be diploid. We assume the population stays at carrying capacity. This implies that juveniles may only settle in replacement of recently deceased adults (Figure 1). Once settled, juveniles reach maturity before the next mating event.

**Figure 1.**
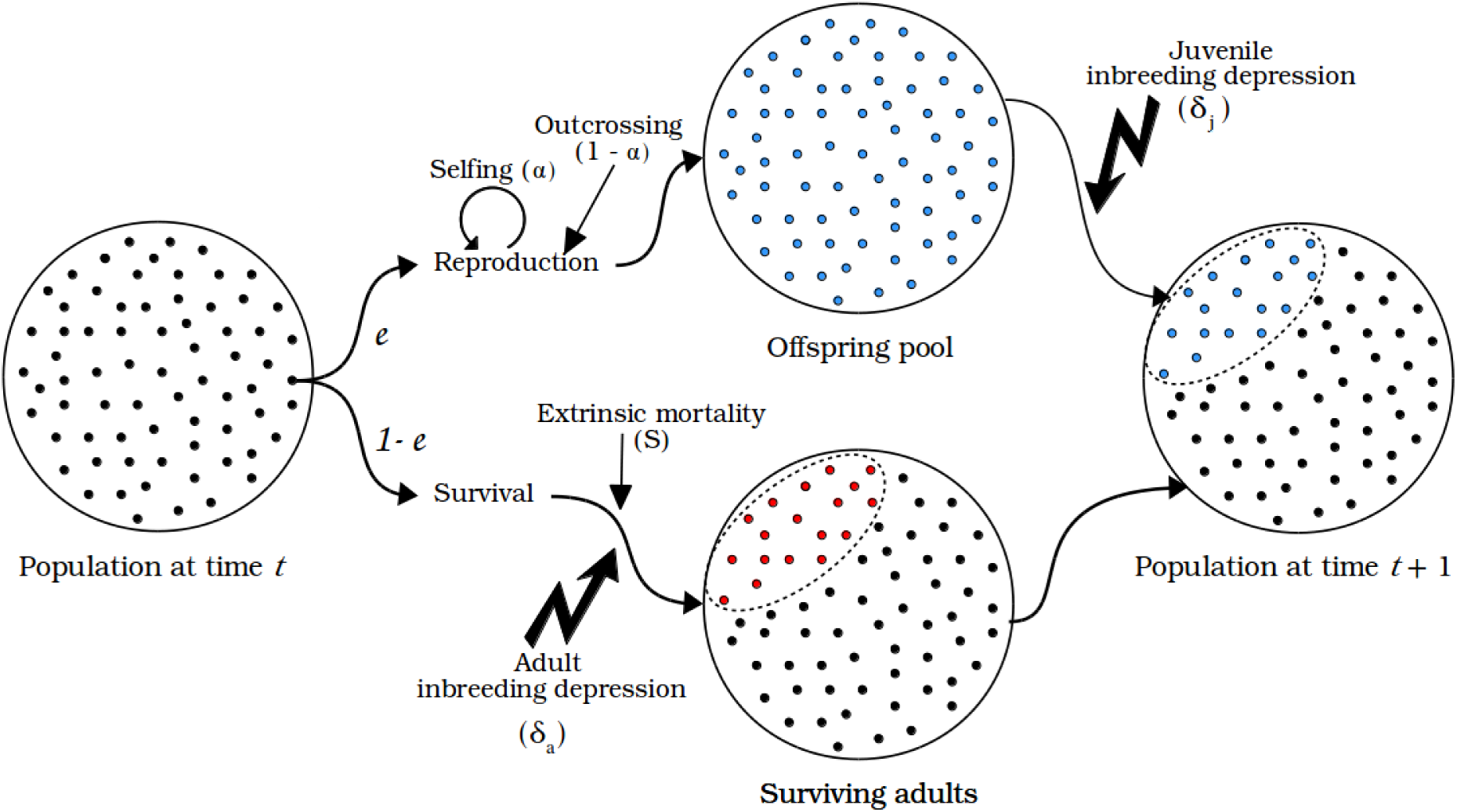
Schematic representation of the life cycle and demography assumed in the model. Established individuals allocate a fraction *e* of their resources to reproduction, and the remaining 1 − *e* to survival. Juveniles replace deceased adults. Red dots depict deceased adults, and blue dots depict juveniles.

We assume that adults keep the same fecundity and survival probability throughout their lives (*i.e.* no age-specific effects). Established individuals allocate a fraction *e* of their resources to reproduction, and the remaining fraction 1 − *e* to post-breeding survival. Consequently, sexually mature individuals have a certain probability of survival between mating events (say, flowering seasons) and generations may overlap: the more resources an individual allocates to reproduction, the larger its reproductive output, but the lower its chances of survival until the next mating event. During each mating event, individuals reproduce by self-fertilisation in a proportion *α*, and by random mating otherwise. Selfed offspring suffer from inbreeding depression [8] differently depending on the considered stage. As juveniles, they suffer from juvenile inbreeding depression, denoted *δ*_*j*_, which decreases their probability of survival to maturity. If they reach maturity, they suffer from adult inbreeding depression, denoted *δ*_*a*_, which decreases their survival probability between mating events. Denoting *𝒮*_*o*_(*e*) the survival probability between two mating events of an outcrossed individual as a function of its reproductive effort, that of a selfed individual, *𝒮*_*s*_(*e*), is therefore given by *𝒮*_*s*_(*e*) = *𝒮*_*o*_(*e*) × (1 − *δ*_*a*_). In simulations, we also considered the case where selfed adults suffer from inbreeding depression on fecundity, which diminishes selfed individuals’ contribution to the gamete pool by a proportion *δ*_*f*_ (Appendix VI).

Whether lifetime inbreeding depression, that is the decrease in lifetime fitness of selfed individuals relative to the outcrossed, varies with life expectancy depends on the life stages we assume inbreeding depression to affect. When inbreeding depression affects juvenile survival or fecundity, lifetime inbreeding depression is unaffected by life expectancy. On the contrary, when inbreeding depression affects adult survival, lifetime inbreeding depression increases with life expectancy (Appendix I). Indeed, in the latter case, selfed individuals have less opportunities to reproduce, while in the former, selfed individuals have the same life expectancy as the outcrossed.

#### Genetic assumptions

We assume that individuals’ selfing rate and reproductive effort are each entirely determined by a single biallelic modifier locus. In each case, we consider the population to be initially fixed with one allele (the resident) and introduce a rare mutant allele which has a small effect on its bearer’s phenotype. We then follow the change in frequency of this mutant, and look for situations where no mutant can increase in frequency, that is, Evolutionarily Stable Strategies [ESS, 22].

### 2.2 Analytical methods

For each model, we obtained analytical predictions for the evolutionary stable strategies, using the theoretical framework introduced by Barton and Turelli [5] and generalized by Kirkpatrick, Johnson, and Barton [19]. Only a summary of the results is given in the main text, detailed recursions can be found in Appendix I, II and III.

### 2.3 Numerical analyses and simulations methods

All programs used in the present study are available on GitHub: https://github.com/Thomas-Lesaffre/M2_project

#### Numerical analyses

The analytical results with approximations we obtain when studying the evolution of reproductive effort and selfing separately are compared with that of exact numerical analyses. For each model, the exact recursions, that is tracking genotypic frequencies (rather than allelic frequencies and genetic associations) with no approximations, are run for 10^6^ generations. A rare mutant with a small effect (*p*_*m*_ = 10^−4^, *ε* = 0.01) is introduced at *t* = 0, and the resulting frequency of the mutant is compared to its initial one. When the mutant increases in frequency, the mutant allele is taken as resident, and recursions are run again introducing a new mutant, until mutants no longer increase in frequency or the analysis hits a bound (*i.e.* 0 or 1). As these analyses were only conducted for validation purpose, outputs are presented in Appendix V. Our results showed that the approximations we obtain are very close to the numerically obtained ESS, when considering the evolution of reproductive effort and selfing separately.

#### Individual-centered simulations

To study the joint evolution of lifespan and selfing, we performed individual-centered C++ simulations, incorporating inbreeding depression affecting juvenile (*δ*_*j*_) and adult survival (*δ*_*a*_). In Appendix VI, we also consider the influence of inbreeding depression affecting fecundity (*δ*_*f*_). In simulations, individuals follow the same life cycle as described above (Fig. 1), and their selfing rate and reproductive effort are each determined by one modifier locus, which are allowed to mutate in both directions, following a uniform distribution in [*α*_0_ - *d, α*_0_ + *d*] and [*e*_0_ - *d, e*_0_ + *d*] where *α*_0_ and *e*_0_ are the parent’s selfing rate and reproductive effort, respectively, with *d* = 10^−1^ and a mutation rate *U*_*m*_ = 10^−2^. Free recombination 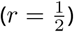 is assumed between the two loci.

## 3. Results

Throughout the following sections, we will need to track the proportion of selfed individuals in the population (Θ), which stays close to its equilibrium value in the absence of mutants (Θ^*\**^) when mutants at modifier loci are rare and only weakly deviate from the resident strategy. As shown in Appendix I, this equilibrium value is given by

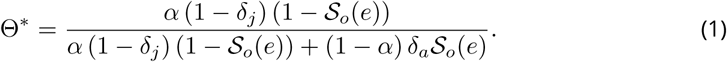

Equation (1) is a decreasing function of *𝒮*_*o*_(*e*), that is the equilibrium proportion of selfed individuals in the population is decreased by overlapping generations. We show in Appendix I that this is due to the repeated effect of adult inbreeding depression on post-breeding survival.

### 3.1 Evolution of lifespan in a partially selfing population

In this section, we analyse a model for the evolution of lifespan under partial selfing through the evolution of reproductive effort, considering the selfing rate *α* as a parameter, and assuming inbreeding depression affects juvenile and adult survival. We assume that the reproductive effort of a given individual is entirely determined by its genotype at a single biallelic modifier locus. Alleles *M* and *m*, which we assume to be codominant, code for a reproductive effort *e* = *e*_0_ and *e* = *e*_0_ + *ε* (*ε ≪* 1), respectively. Furthermore, following Zhang [33], we assume that the survival probability between two mating events of an outcrossed individual as a function of its reproductive effort *e*, is given by

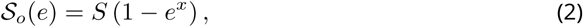

where *S* is the maximal survival probability, that is a measure of extrinsic mortality, and *x* controls the shape of the survival *vs* reproduction trade-off. We use this function form because it is flexible and allows for the consideration of a variety of trade-off shapes. The detailed recursions are given in Appendix II. In brief, we follow the variation of three variables, namely, the frequency of allele *m* (*p*_*m*_), which is assumed to be a very rare mutant with a small effect (*ε* ≪ 1), the deviation in homozygosity at the modifier locus as compared to the panmictic expectation (*D*_*m,m*_), and the proportion of selfed individuals in the population (Θ), which is assumed to remain close to its equilibrium value Θ^*\**^. We look for Evolutionarily Stable Strategies [ESS, 22], that is situations where no mutant allele *m* may invade the population and replace the resident allele *M*.

#### Allelic frequencies change

To leading order in *ε*, we can express the change in frequency of allele *m* between two timesteps (Δ*p*_*m*_) as

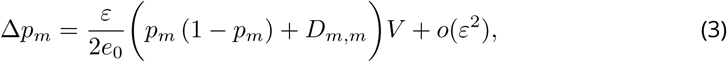

with *V* = 1 − *S* (1 − Θ*δ*_*a*_) (1 + (*x -* 1) *e*_0_^*x*^). Using a separation of timescales approximation [19], that is assuming the deviation in homozygosity at the modifier rapidly equilibrates in comparison with allelic frequencies, we obtain a quasi-equilibrium value by solving Δ*D*_*m,m*_ = 0 for *D*_*m,m*_ (where Δ*D*_*m,m*_ is the change in excess in homozygotes at the modifier between two timesteps). Since *D*_*m,m*_ ≥ 0 provided that *ε ≪* 1, the first term in Equation (3) is always of the sign of *ε*, and only *V* matters for the determination of the equilibrium.

#### Evolutionarily Stable Strategy

In order to determine the equilibrium reproductive effort *e*^*\**^, we plug Equation (1) into *V,* and solve *V* = 0 for *e*_0_, which yields

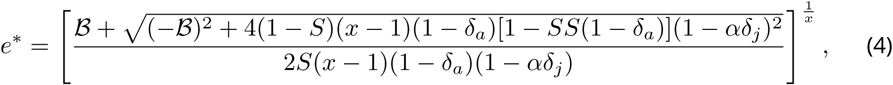

with *ℬ* = (1 − *S*)(2 - *x*)(1 −α *δ*_*j*_) -*δ* _*a*_ [(1 − *S*(2 - *x*))(1 − α *δ*_*j*_) - *x*α (1 − *δ*_*j*_)].

For this equilibrium to be stable, that is for Equation (4) to be an ESS, it is required that

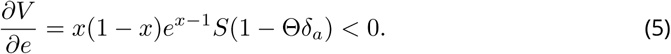

One can see that when the survival *vs.* reproduction trade-off is convex or linear (*x* ≼ 1), condition (5) is not fulfilled, which implies that Δ*p*_*m*_ *≽* 0. Thus, alleles increasing their bearer’s reproductive effort are always favoured, and only annuality (*e*^*\**^ = 1) can evolve. On the other hand, when the trade-off is concave 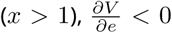, Equation (4) is an ESS and perennial strategies can be maintained. This result is classical, and has been reached by numerous authors before [e.g. 6, 27, 33]. In addition, Equation (4) is a decreasing function of *S*. That is, higher extrinsic mortality favours higher reproductive efforts: allocating resources to survival becomes less and less advantageous as extrinsic mortality increases (*i.e.* as *S* decreases), because individuals are more likely to die independently of the resources they spend on survival. Hence, our model also predicts the classical observation that annuals are more likely to evolve in more disturbed habitats, where extrinsic mortality is higher.

Assuming *x* > 1, that is a concave trade-off, differentiating Equation (4) with respect to *α* yields

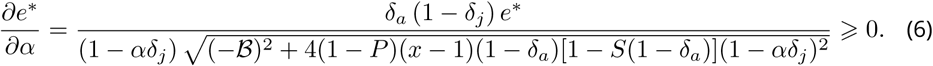

Equation (6) shows that the ESS reproductive effort is an increasing function of the selfing rate *α*, as long as some selfed juveniles survive to maturity (*δ*_*j*_ ≠ 1), and adults suffer from inbreeding depression (*δ*_*a*_ ≠ 0). In other words, through its impact on adult survival, self-fertilisation favours evolution towards higher reproductive efforts, and hence shorter lifespans. This effect becomes stronger as adult inbreeding depression increases, because allocating resources to survival becomes less beneficial in selfed adults, and as juvenile inbreeding depression decreases, because it reduces the proportion of selfed individuals entering the adult population. Therefore, we show that self-fertilisation favours evolution of shorter lifespans.

### 3.2 Influence of lifespan on the evolution of self-fertilisation

We now study the influence of lifespan on the evolution of self-fertilisation assuming inbreeding depression only affects juvenile and adult survival, taking reproductive effort, *i.e.* lifespan, as a fixed parameter. We assume that the selfing rate of a given parent is entirely determined by a single biallelic locus. Alleles *M* and *m*, which are assumed to be codominant, code for selfing rates *α*_0_ and *α*_0_ + *a,* respectively. Allele *m* is assumed to be a rare mutant with a weak effect (*a* ≪ 1). In this section, we do not need to make any assumption regarding the shape of the survival *vs* reproduction trade-off. Hence, the general *𝒮* _*o*_(*e*) notation will be used. Recursions are given in Appendix III. As in the previous section, we follow three variables: the frequency of the mutant allele (*p*_*m*_), the deviation in homozygosity at the modifier locus as compared to the panmictic expectation (*D*_*m,m*_), and the proportion of selfed individuals in the population (Θ), which is assumed to remain close to its equilibrium value (Θ^*\**^, Equation (1)) when the mutant is rare. We look for the conditions under which allele *m* invades the population.

#### Allelic frequencies change

Plugging Equation (1) into the allelic frequencies change (Δ*p*_*m*_) yields, to leading order *a*,

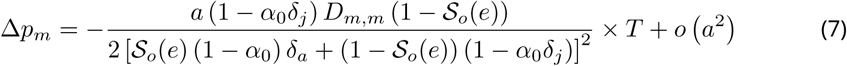

with *T* = [1-*𝒮*_*o*_(*e*) (1 + *δ* _*a*_] [*𝒮*_*o*_(*e*) *δ* _*a*_*-*(1-*𝒮*_*o*_(*e*)) (1 − 2*δ* _*j*_)].

Using the same arguments as in the previous section, we show in Appendix III that for all *α, D*_*m,m*_ > 0. Hence, the first term in Equation (7) is always negative, and only *T* matters for the determination of the equilibrium. Since *T* does not depend on the selfing rate, there are only two possible situations: either *T* > 0 and full outcrossing (*α* = 0) is favoured, or *T* < 0 and full selfing evolves (*α* = 1), similar to the findings of previous authors [e.g. 20]. Solving *T* < 0 for *δ*_*j*_, we have

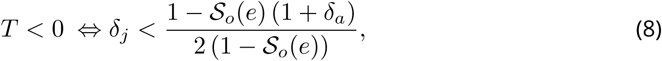

which simplifies to *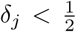* in the annual case (*𝒮*_*o*_(*e*) = 0). Otherwise, this threshold is a decreasing function of *𝒮*_*o*_(*e*) when *δ*_*a*_ > 0. Hence, we find that the range of conditions under which self-fertilisation can evolve in a population decreases when lifespan increases, provided that inbreeding depression affects adult survival, in agreement with previous results [24]. This result implies that even very weak adult inbreeding depression is sufficient to prevent evolution of self-fertilisation in long-lived species. Equation (8) was validated using numerical analyses, which were found to be in very good agreement with the analytical prediction (Appendix V).

### 3.3 Joint evolution of lifespan and self-fertilisation

In this final section, we study the joint evolution of lifespan and selfing. Figure 2 highlights the different situations that arise when considering the joint evolution of lifespan and selfing. The threshold reproductive effort above which selfing can evolve (*ē*) can be obtained by setting *𝒮*_*o*_(*e*) = *S* (1 − *e*^*x*^) in Equation (8), and solving *T* < 0 for *e*. It is given by

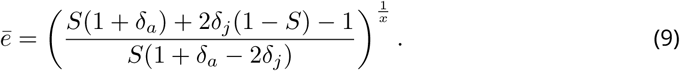

**Figure 2.**
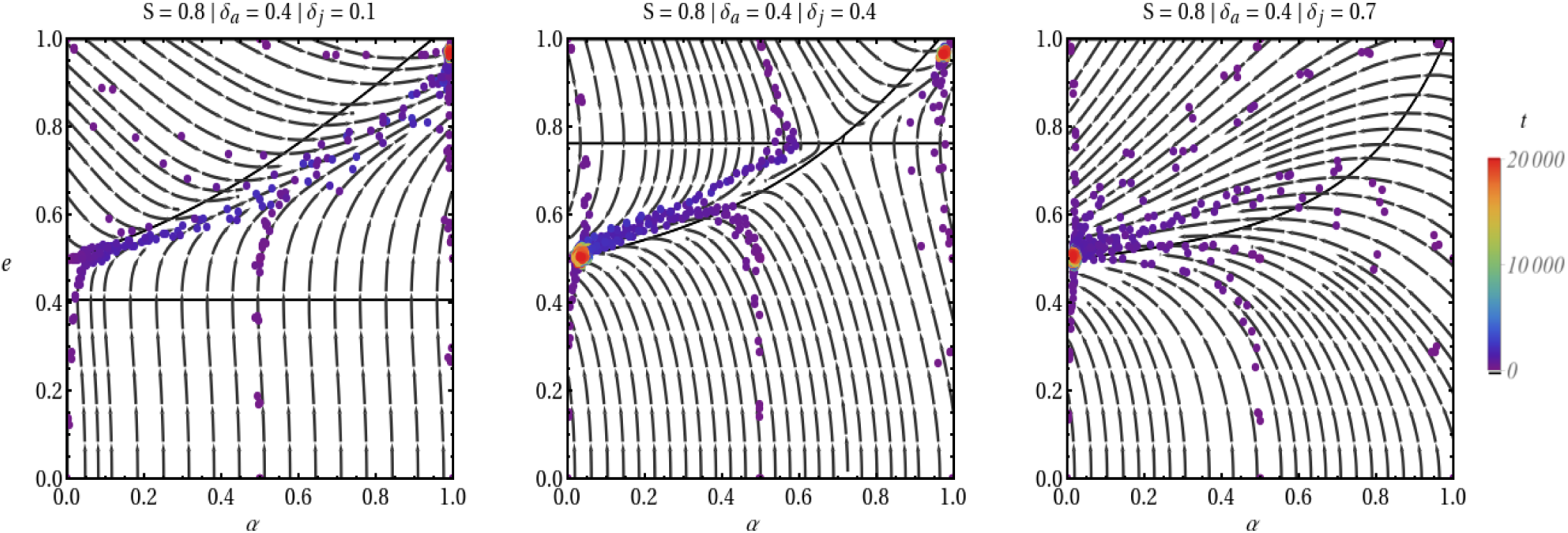
Phase diagrams highlighting the three kinds of behaviour that can arise from the coevolutionary dynamics of lifespan and selfing. Reproductive effort (*e*) is plotted against selfing rate (*α*). Solid lines depict isoclines, and arrows indicate how the joint evolution behaves. Points depict simulation results, and are colored with respect to time.

Overall, the transition to selfing is always associated with increased reproductive effort, that is reduced lifespan. Depending on the inbreeding depression and extrinsic mortality conditions, one of three things can happen: (1) selfing is always favored (Fig. 2a), and reproductive effort evolves to

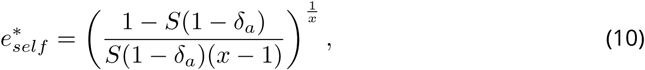

(2) outcrossing is always maintained (Fig. 2c), and reproductive effort stabilises at

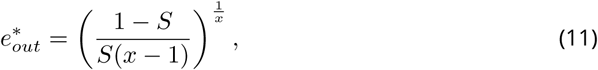

(3) either of the two stable equilibria (*α* = 0 and *α* = 1) can be reached, depending on initial conditions (Fig. 2b). This occurs when adult inbreeding depression is high enough, provided that *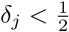* (Fig. 3). Selfers then stabilise at a high reproductive effort, which allows them to escape the deleterious effects of adult inbreeding depression. In other words, in situations where extrinsic mortality conditions would allow for the evolution of longer lifespans, selfers could remain short-lived solely owing to the deleterious effects of inbreeding.

**Figure 3.**
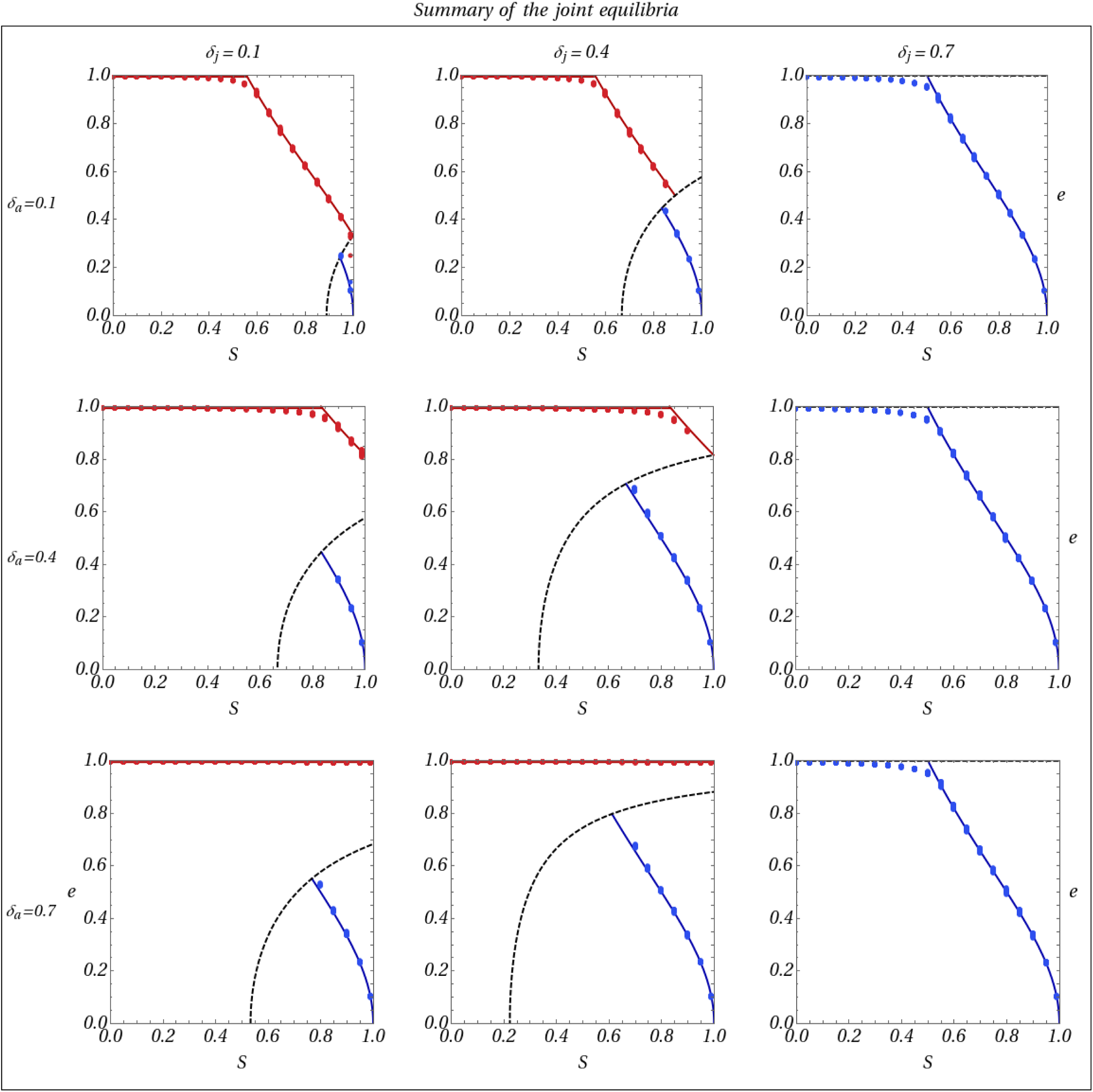
Joint evolutionary equilibria. All results presented here assume *x* = 2. Y-axis is the ESS reproductive effort (*e*^*\**^), and X-axis is extrinsic mortality (*S*). The lines correspond to analytical results, while dots depict simulations results. The red ones corresponds to the equilibrium reproductive effort when *α* = 1 is reached, while the blue line corresponds to the equilibrium reproductive effort when *α* = 0 is reached. The dashed black line represents the threshold lifetime inbreeding depression under which outcrossing can be maintained.

Figure 3 summarizes the evolutionary stable outcomes of the joint evolution of lifespan and selfing in various inbreeding depression and extrinsic mortality conditions. Simulation results are in good agreement with analytical predictions, although small deviations are observed in Figure 2 and 3 for high and low selfing rates and reproductive efforts. These deviations are a consequence of the value constraints applied to the traits to keep them within bounds. Indeed, as the trait value (*i.e.* selfing rate or reproductive effort) approaches a bound of its definition domain (that is, 0 or 1 on the [0, 1] interval), it will tend to mutate slightly away from it, as all mutations exceeding the bound will be cut back to the bound value (for example, a mutation at the reproductive effort modifier coding for a reproductive effort *e* > 1 will be cut to *e* = 1).

In Appendix VI, we also consider the effect of inbreeding depression affecting fecundity (*δ*_*f*_). We show that *δ*_*f*_ or *δ*_*j*_ have the same effect: they reduce the range of conditions under which the full selfing strategy can evolve. The mechanisms underlying these similar behaviours are however slightly different. Indeed, while in both cases they are caused by a reduction of the participation of selfed individuals to reproduction, *δ*_*j*_ decreases the proportion of selfed individuals in the population. On the other hand, *δ*_*f*_ reduces the contribution of selfed adults to reproduction by reducing their contribution to the gamete pool.

## 4 Discussion

In this paper, we built modifier models to investigate the joint evolution of lifespan and mating system. To the best of our knowledge, this is the first time modifier models are used to study the evolution of life-history traits in a population with overlapping generations.

We found only two stable mating systems, full outcrossing and full selfing. In agreement with previous authors, we found that increasing lifespan considerably reduces the range of conditions under which self-fertilisation can evolve, because of the repeated effect of adult inbreeding depression which results in high lifetime inbreeding depression [24]. Furthermore, we found inbreeding depression on juvenile survival and fecundity (Appendix VI) to affect the coevolutionary dynamics similarly, by reducing the range of conditions under which the full selfing strategy can be reached. Conversely, we showed that self-fertilisation is expected to cause evolution towards shorter lifespans due to inbreeding depression affecting adult survival, even under the low inbreeding depression conditions required for selfing to evolve in long-lived species. In addition, we showed that self-fertilising species could remain short-lived even in a very stable habitat, as a strategy to avoid the deleterious effects of inbreeding. Finally, we showed that long-lived selfers can only emerge under very weak adult inbreeding depression. Overall, our results agree with the well-documented empirical association of short lifespans with selfing, and long lifespans with outcrossing [25].

The only previous study of the influence of the mating system on the evolution of lifespan is that of Zhang [33]. While our main conclusions are in line with theirs, they also differ in a number of ways. Zhang [33] concluded that self-fertilisation could induce evolution of shorter lifespans through increased allocation to female reproduction, without any role for adult inbreeding depression, provided that female reproduction is sufficiently costly, that inbreeding depression is low and that juvenile survivorship is constrained within a narrow range of values. Zhang [33]’s results and ours are not mutually exclusive, since they rely on different mechanisms. However, we expect our prediction to apply more generally, as it only requires inbreeding depression to affect adult survival to apply, which seems reasonable, as inbreeding depression commonly occurs over all stages of life [17].

Zhang [33]’s approach and ours also differ widely in the methods used. While in our model, investing more or less resources into reproduction directly affects the probability to remain in the population, and therefore to carry out more mating events, Zhang [33] modeled adult survival as an additive term (see Equation (3) in their study). In our opinion, this way of modelling survival does not incorporate the fact that surviving not only allows to bring one’s own gene copies to the next flowering season, but also to carry out more mating events, and therefore merely depicts survival as a poor alternative to reproduction for gene transmission to the next flowering season. This implies that the survival *vs.* reproduction trade-off is modelled incorrectly. In Appendix IV, we present an alternative approach based on a phenotypic model, by which conclusions similar to those we reach in the present study can be obtained.

Overall, both theoretical and empirical results indicate that predominantly selfing species are almost always short-lived, while predominantly outcrossing ones are mostly long-lived. It is difficult to identify the causative mechanisms underlying such a correlation. The role of models such as the present study is to provide putative ecological and genetic conditions under which a given association may emerge. Given these results, we propose that lifespan could act as a confounding factor when considering the joint evolution of self-fertilisation with other traits. For example, lifespan shortening following the transition to self-fertilisation could in part be responsible for the emergence of the selfing syndrome, rather than self-fertilisation *per se*. Indeed, as a shorter lifespan implies that individuals have to complete their cycle more rapidly, they could grow smaller flowers regardless of an adaptation to more efficient selfing [29].

Accounting for lifespan could also shed new light on some long-standing evolutionary questions related to the evolution of mating systems. For instance, lifespan could influence the joint evolution of dispersal and self-fertilisation in two ways. First, increased lifespan induces higher local relatedness [13], and perennials tend to occupy more saturated, competitive environments. Hence, kin competition should be greater in more long-lived species, and long-distance dispersal could be favoured as a mean to avoid it. Since perennials outcross more than annuals, this could generate an indirect association between long-distance dispersal and outcrossing through lifespan. Second, the repeated effect of adult inbreeding depression considerably increases the consequences of mating among relatives in perennials. Long-distance dispersal and outcrossing could thus be favoured jointly in such species as an inbreeding avoidance strategy [3].

Throughout this work, we assumed fixed inbreeding depression. Inbreeding depression is generally thought to be caused by recessive deleterious mutations segregating at low frequencies in populations [9]. The population genetics of such mutations in populations with overlapping generations are poorly understood theoretically. Evidence from large-scale meta-analyses suggest that the measured magnitude of inbreeding depression increases as species’ lifespan increases [2, 12]. However, it is unclear whether this pattern is due to mating system differences between long-lived and short-lived species [25], or to lifespan. Additionally, empirical studies of inbreeding depression in perennials are rather scarce, and rarely span over several years, let alone individuals’ entire lives [17]. Consequently, the quantities measured in such studies are not likely to depict inbreeding depression in its classic definition, that is the lifetime fitness decrease of selfed individuals as compared to outcrossed ones [8], but rather its magnitude at a given timestep or stage. Therefore, estimates in short-lived and long-lived species may not be readily comparable, and higher inbreeding depression levels reported in perennials could in part reflect measurement biases [2]. Lifespan may also interact with the mutation load in non-trivial ways. On the one hand, longer lifespan may increase selection against deleterious mutations, because of the increased number of opportunities for selection to occur, thereby leading to lower levels of inbreeding depression through better purging [23]. On the other hand, more long-lived species may endure significantly more mitotic mutations throughout their lives owing to their overall larger stature, which could result in an increase in inbreeding depression as plants do not have a separated germline [28]. Furthermore, perennials may experience reduced purging of mutations affecting juvenile fitness, as some of these mutations would remain as neutral mutations among adults, and be recurrently reintroduced in the population through reproduction [1].

Importantly, these various predictions stem from vastly different approaches, and remain mostly verbal. In particular, the rare models that considered populations with overlapping generations vary considerably in terms of the life stage they assume deleterious mutations to affect. Indeed, while some authors assume they only act on juvenile survival or gametes production [1], others assume they affect their bearers’ survival throughout their lives [23]. As for the accumulation and transmission of somatic mutations [28], the population-level consequences of this mechanism were never investigated theoretically. Besides, no study has yet considered the interaction between separate loads affecting different life stages, although these situations are likely to occur since differential purging has been reported between life stages [2]. Finally, every theoretical study so far has assumed individuals’ fecundities and survivorship to not depend upon their age, which is a strong simplifying assumption [15, 26]. In the future, significant insight for the evolution of mating systems and life-histories is to be gained through investigations of the dynamics of mutation loads in perennials.

## Acknowledgements

This preprint has been reviewed and recommended by Peer Community In Evolutionary Biology (https://dx.doi.org/10.24072/pci.evolbiol.100070). We thank Thomas Bataillon, along with two anonymous reviewers, for taking the time to consider our manuscript, and ultimately recommending our work. Their comments greatly improved the quality of the manuscript. We also thank Sylvain Glémin, Florence Débarre, Diala Abu Awad, Vincent Castric, Louis Mahé and Denis Roze for helpful discussions and comments on the manuscript. This work was funded by the European Research Council (NOVEL project, grant #648321). The authors also thank the Région Hauts-de-France, and the Ministère de l’Enseignement Supérieur et de la Recherche (CPER Climibio), and the European Fund for Regional Economic Development for their financial support.

## Conflict of interest disclosure

The authors of this preprint declare that they have no financial conflict of interest with the content of this article. SB is one of the PCI Evol Biol recommenders.

## APPENDIX I

### Proportion of selfed individuals and lifetime inbreeding depression

In this section, we detail the derivation of the equilibrium proportion of selfed individuals in the population, and show that its decrease when longevity increases is attributable to stronger selection against inbred individuals.

#### Proportion of selfed individuals

Following reproduction, adult individuals survive with probability *𝒮*_*o*_(*e*) if they were produced by outcrossing, and (1 − *δ*_*a*_) *𝒮*_*o*_(*e*) if they were selfed. Therefore, denoting Θ the proportion of selfed individuals in the population at a given timestep, the proportion of adult individuals surviving between two mating events 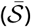 is given by

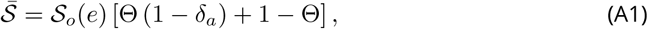

and the proportion of selfed individuals among them, Θ^*s*^ is

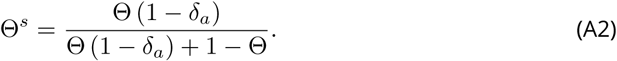

Among the juveniles settling in the population at a given mating event, the proportion of selfed individuals (Θ^*j*^) given the selfing rate *α* can be expressed as

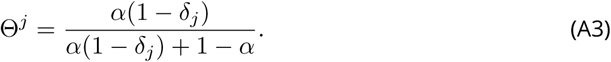

Hence, the proportion of selfed individuals at the next timestep (Θ*′*) is

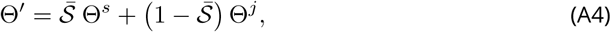

and, solving ΔΘ = Θ*′* − Θ = 0, we obtain the equilibrium proportion of selfed individuals in the population, Θ^*\**^:

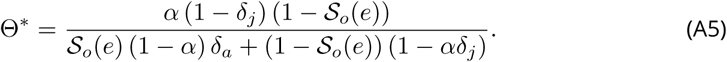

#### Lifetime inbreeding depression

In our model, inbreeding depression only affects survival, in juveniles and in adults. Thus, the decrease in fitness selfed individuals suffer from in comparison with the outcrossed throughout their lives, and not only between two mating events (*i.e.* lifetime inbreeding depression) can be expressed in terms of life expectancies. The life expectancies for selfed (*ℒ*_*s*_) and outcrossed (*ℒ*_*o*_) individuals, respectively, are given by

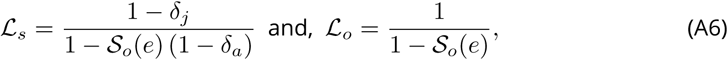

and lifetime inbreeding depression can be expressed as

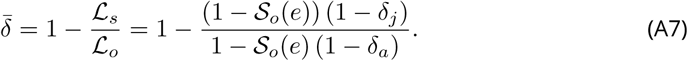

Equation (A7) is an increasing function of *𝒮* _*o*_(*e*), that is selection against selfed individuals grows stronger as longevity increases. This is due to the repeated effect of adult inbreeding depression on post-breeding survival.In the annual case (*𝒮* _*o*_(*e*) = 0), Equation (A7) reduces to *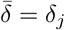* Interestingly, Equation (A7) relates to Θ^*\**^ (Equation (A5)) in the following way:

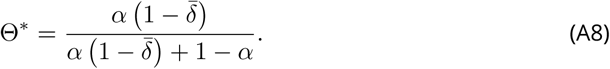

Therefore, the equilibrium proportion of selfed individuals in the population decreases when generations overlap (that is, *𝒮* _*o*_(*e*)) increases, due to stronger selection against selfed individuals owing to the repeated effect of adult inbreeding depression on adult survival.

## APPENDIX II

### Recursions for the evolution of reproductive effort in a partially selfing population

In this section, we detail the mathematical derivations leading to the ESS reproductive effort expression we obtain. The selfing rate is fixed and reproductive effort is controlled by a single modifier locus. To describe the changes happening in the population between two timesteps, we use the theoretical framework first described by Barton and Turelli [5] and generalized by Kirkpatrick, Johnson, and Barton [19]. We follow the variation of three variables, namely, the frequency (*p*_*m*_) of the mutant allele, which is assumed to be a rare with a small effect (*∈*) on its bearer’s reproductive effort, the excess in homozygosity at the modifier locus (*D*_*m,m*_), and the proportion of selfed individuals in the population (Θ).

We define the indicator variables *X*_*m*_ and *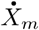*, corresponding to the two allelic positions of the modifier locus located on the paternally and maternally inherited chromosomes, respectively. These variables can take two values, 1 if the mutant allele (*m*) is present at the considered position, and 0 otherwise. Since there is no maternal or paternal effect on the expression of the alleles at the modifier, we have:

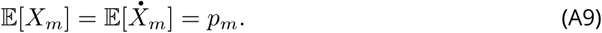

Using these indicator variables, we may define the centered variables *ζ*_*m*_ and *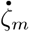* as:

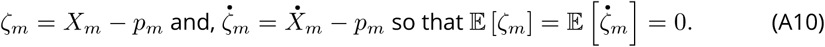

These variables allow us to define the excess in homozygotes in the population at the modifier as compared to the panmictic expectation (*D*_*m,m*_) as:

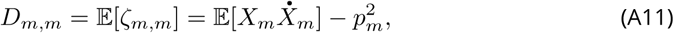

Where 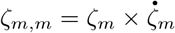.

#### Reproduction

Using indicator variables, the reproductive effort of a given individual, *e*, can be expressed as follows

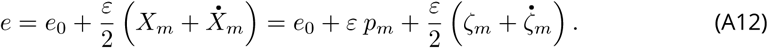

The average reproductive effort is thus

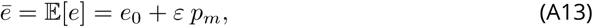

which yields, to leading order in *ε*, the relative contribution of a given individual to reproduction 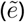 during a given mating event

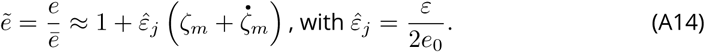

#### Frequency of the mutant

Using Equation (A14), the frequency of the mutant allele 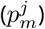 among juveniles following reproduction is

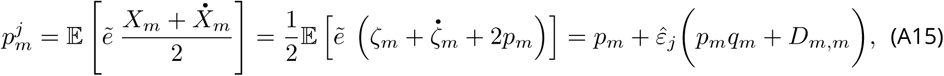

with *q*_*m*_ = 1 − *p*_*m*_, and the change in frequency of the mutant among juveniles is therefore

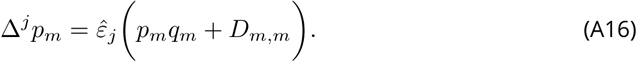

#### Excess in homozygotes

The change in homozygosity at the modifier in juveniles can be divided in two phases, selection and syngamy. The excess in homozygotes at the modifier following selection, *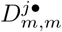* is given by

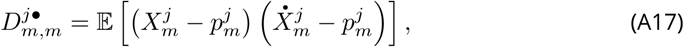

where *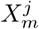* and 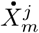 are indicator variables defined among juveniles following selection (so that 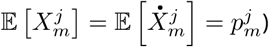. Noting that *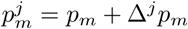*, and expressing Equation (A17) in terms of *ζ*-variables, we have

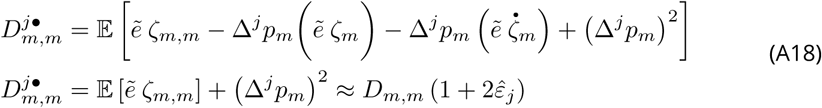

to leading order in *ε* and assuming *p*_*m*_ is small. During syngamy, homozygosity is generated by inbreeding. In our model, we assumed partial selfing. Therefore, the excess in homozygotes among juveniles, after accounting for juvenile inbreeding depression, is given by

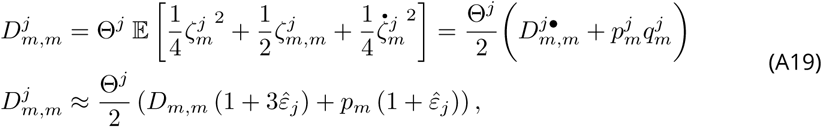

where 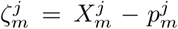 (resp.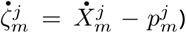 is the centered variable defined for the paternally (resp. maternally) inherited chromosome among juveniles following selection, that is taking the frequency of the mutant among juveniles following selection 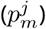 as reference value [19].

#### Survival

We denote by *𝒮* the probability of survival of a parent taken at random in the parental population. Using our indicator variables, and given the proportion Θ of selfed individuals in the population, the probability of survival is

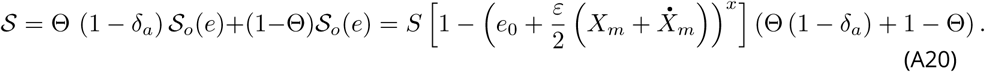

Substituting indicator variables with *ζ*-variables and assuming that *∈* and *p*_*m*_ are small, we obtain, to leading order,

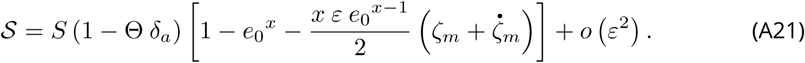

Thus, the mean survival probability is

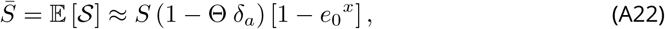

and the relative survival probability 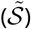 simplifies to

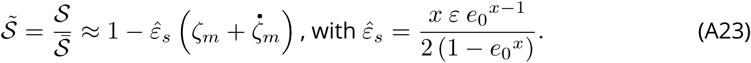

#### Frequency of the mutant

Using Equation (A23), the frequency of the mutant 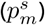 among the surviving parents is

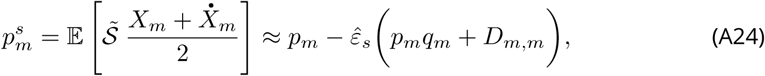

and the variation in frequency of the mutant owing to selection among parents is therefore given by

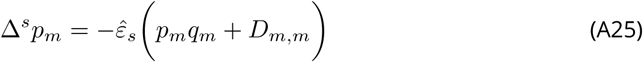

#### Excess in homozygotes

Since only selection acts at this stage, the excess in homozygotes at the modifier locus among the parents 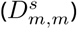 is given by

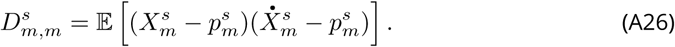

Noting that *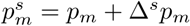*, and expanding Equation (A26), we have

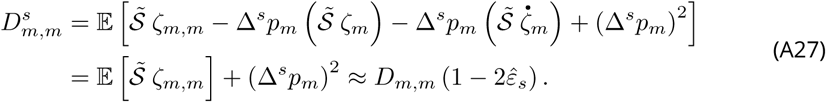

#### Next timestep

From our previously derived expressions, the frequency of the mutant 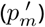, the excess in homozygotes 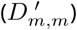 and the proportion of selfed individuals in the population (Θ*′*) in the next timestep is

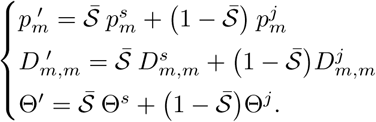

In order to determine the *Evolutionary Stable Strategy* [ESS, 22] for the population given the parameters of the model, one has to determine the value of *e*_0_ for which no mutant allele (*m*) may invade the population and replace the resident (*M*). To do so, we will consider the change in frequency of the mutant allele over one timestep (Δ*p*_*m*_), which can be expressed as

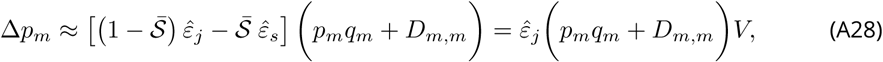

with *V* = 1 − *S* (1 – Θ*δ* _*α*_) (1 + (*x* - 1) *e*_0_^*x*^).

#### Separation of timescales approximation

When selection is weak (*ε* is small), we may assume that the excess in homozygotes in the population reaches a value close to equilibrium much faster than the allelic frequencies. It is obtained by solving

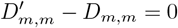

for *D*_*m,m*_. Thus, assuming the mutant is rare (*p*_*m*_ is of order *ε*) we have

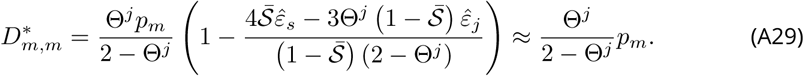

Since we have *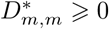* (Equation (A29)), the first term in Equation (A28) is always positive, and only *V* matters for the determination of the equilibrium.

#### Evolutionarily Stable Reproductive Effort

Injecting Equation (A5) into Equation (A28), and solving Δ*p*_*m*_ = 0 for *e*_0_ yields the following ESS for the reproductive effort (*e*^*\**^):

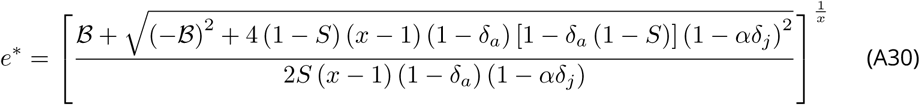

with *B* = (1 − *S*) (2 - *x*) (1 − *αδ*_*j*_) + *δ*_*a*_ [*αx* (1 − *δ*_*j*_) *-* [1 − *S* (2 - *x*)] (1 − *αδ*_*j*_)].

## APPENDIX III

### Recursions for the evolution of self-fertilisation in a perennial population

In this section, we detail the mathematical derivations for the evolution of self-fertilisation in a perennial population. Individuals reproduce by self-fertilisation at a rate given by their genotype at a single biallelic modifier locus, and by random mating otherwise. At the modifier, alleles *M* and *m* are assumed to be codominant, and code for selfing rates *α*_0_ and *α*_0_ + *a*, respectively. Allele *m* is assumed to be a rare mutant with a weak effect (*i.e. a* is small). The theoretical framework we use is the same as in Appendix II.

#### Reproduction

Let us split the population into two groups: the *selfed* and the *outcrossed* individuals. The frequency of allele *m* in each of these groups are denoted *p*_*m,s*_ and *p*_*m,o*_, respectively, so that its frequency in the whole population is given by

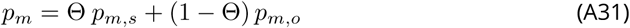

where Θ is the proportion of selfed individuals in the population.

#### Frequency of the mutant among the selfed

The contribution of a given parent to the production of selfed offspring, *α*, is

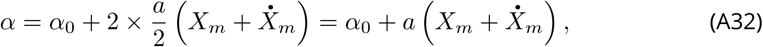

owing to the fact that increasing its selfing rate by 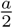 induces a twofold transmission advantage in comparison with others [**fisher1941**]. Injecting *ζ*-variables into Equation (A32), we obtain the following expression for the relative contribution of a given parent to the production of offspring by self-fertilisation:

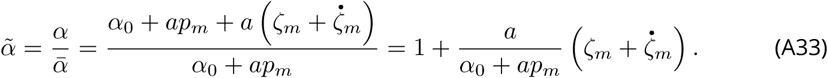

Thus, the frequency of allele *m* among the selfed offspring 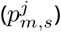 is given by

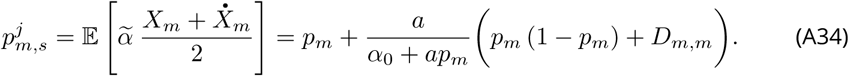

#### Frequency of the mutant among the outcrossed

The contribution of a given parent to the production of outcrossed offspring, *o*, is

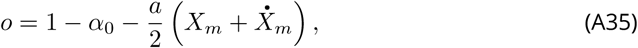

which yields the following expression for its relative contribution (*Õ*):

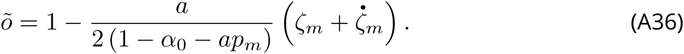

Hence, the frequency of allele *m* among the outcrossed offspring 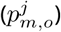 is

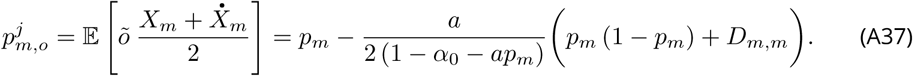

#### Excess in homozygotes

Considering the whole offspring pool, we may neglect the effect of the mutant on the selfing rate. Thus, we may express the excess in homozygotes at the modifier among juveniles as

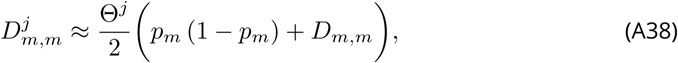

where Θ^*j*^ is the proportion of selfed individuals among the juveniles settling in the population (Equation (A3)).

#### Survival

Among the parents, allelic frequencies among the selfed and among the outcrossed do not vary, because there is no direct selection acting on the modifier among them. Therefore,

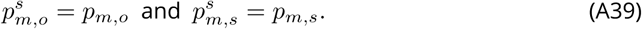

Moreover, we have

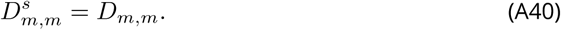

Selfed individuals are however counterselected due to adult inbreeding depression. Thus, their proportion varies according to Equation (A2).

#### Next timestep

In the next timestep, assuming the proportion of selfed individuals remains close to its equilibrium value (Θ^*\**^, Equation (A5)), the frequency of the mutant 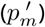 and the excess in homozygotes 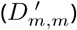 are given by

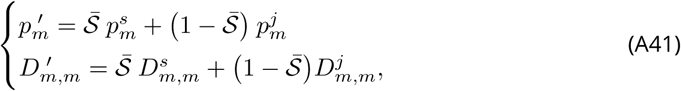

where *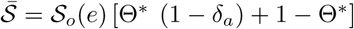* is the proportion of parents surviving until the next timestep. Using these recursions, we may express the change in allelic frequencies at the modifier (Δ*p*_*m*_) as

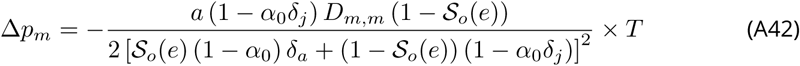

with *T* = [1 − *𝒮*_*o*_(*e*) (1 + *δ*_*a*_)] [*𝒮*_*o*_(*e*)*δ*_*a*_ *-* (1 − *𝒮*_*o*_(*e*)) (1 *-* 2*δ*_*j*_)].

#### Separation of timescales approximation

Assuming homozygosity equilibrates quickly in comparison with allelic frequencies at the modifier, we obtain a quasi-equilibrium expression [19] for the excess in homozygotes at the modifier 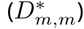 by solving *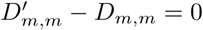*,

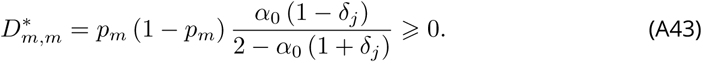

Using Equation (A43), one can see that the first term in Equation (A42) is always negative, and only *T* matters for the determination of the equilibria. Because *T* does not depend on *α*_0_, there is only two situations: either *T* > 0 and full outcrossing is maintained (*α*^*\**^ = 0), or *T* < 0 and full selfing is favoured (*α*^*\**^ = 1). Hence, by solving *T* < 0, we obtain the following threshold for the evolution of self-fertilisation:

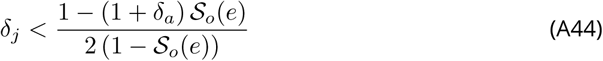

## APPENDIX IV

### Phenotypic approach to the evolution of self-fertilisation and the evolution of lifespan

In this section, we derive a phenotypic approach to the evolution of self-fertilisation in a perennial population, and to the evolution of longevity in a partially selfing population. We express the fitness of the mutant as its *lifetime* reproductive success.

#### Evolution of longevity

All individuals have the same reproductive effort *e* during each mating event they participate in. Therefore, an individual’s fitness (*W*), given its life expectancy (*ℒ*), is given by

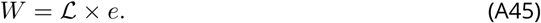

With probability *α*, an individual is produced by self-fertilisation and its life expectancy (*ℒ*_*s*_) is

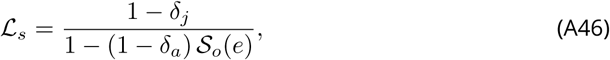

while with probability 1 − *α*, it is outcrossed, and its life expectancy (*ℒ*_*o*_) is given by

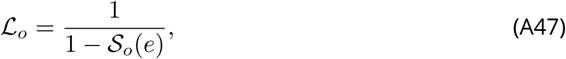

with *𝒮*_*o*_(*e*) = *𝒮* (1 − *e*^*x*^). Therefore, we have

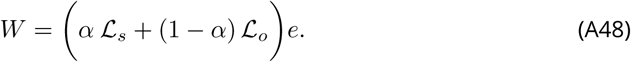

Deriving Equation (A48) once with respect to *e*, we obtain

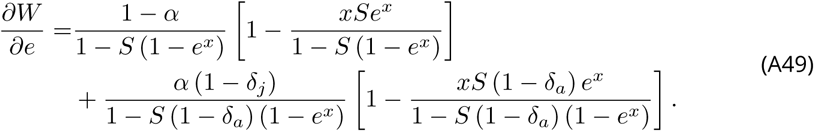

Finally, solving 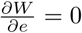 for *e*, we obtain a high order polynomial expression for the equilibrium reproductive effort. In cases where *α* = 0 and *α* = 1, it reduces to

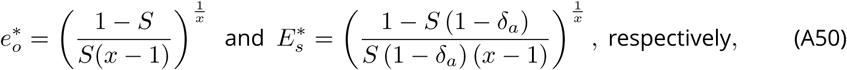

which corresponds to Equations (11) and (10). Although its graphical behaviour is similar to that of the prediction we obtained in our main model (Figure A1, the analytical properties of this solution cannot be studied because of its complexity. Its stability cannot be easily studied analytically either, because the second derivative of the fitness with respect to *e* is too complicated. It can however be assessed graphically. This highlights the interest of using modifier models rather than a classical phenotypic approach for the study of populations with overlapping generations: it allows one to neglect inconsequential terms in order to obtain understandable solutions.

#### Evolution of self-fertilisation

**Figure A1.**
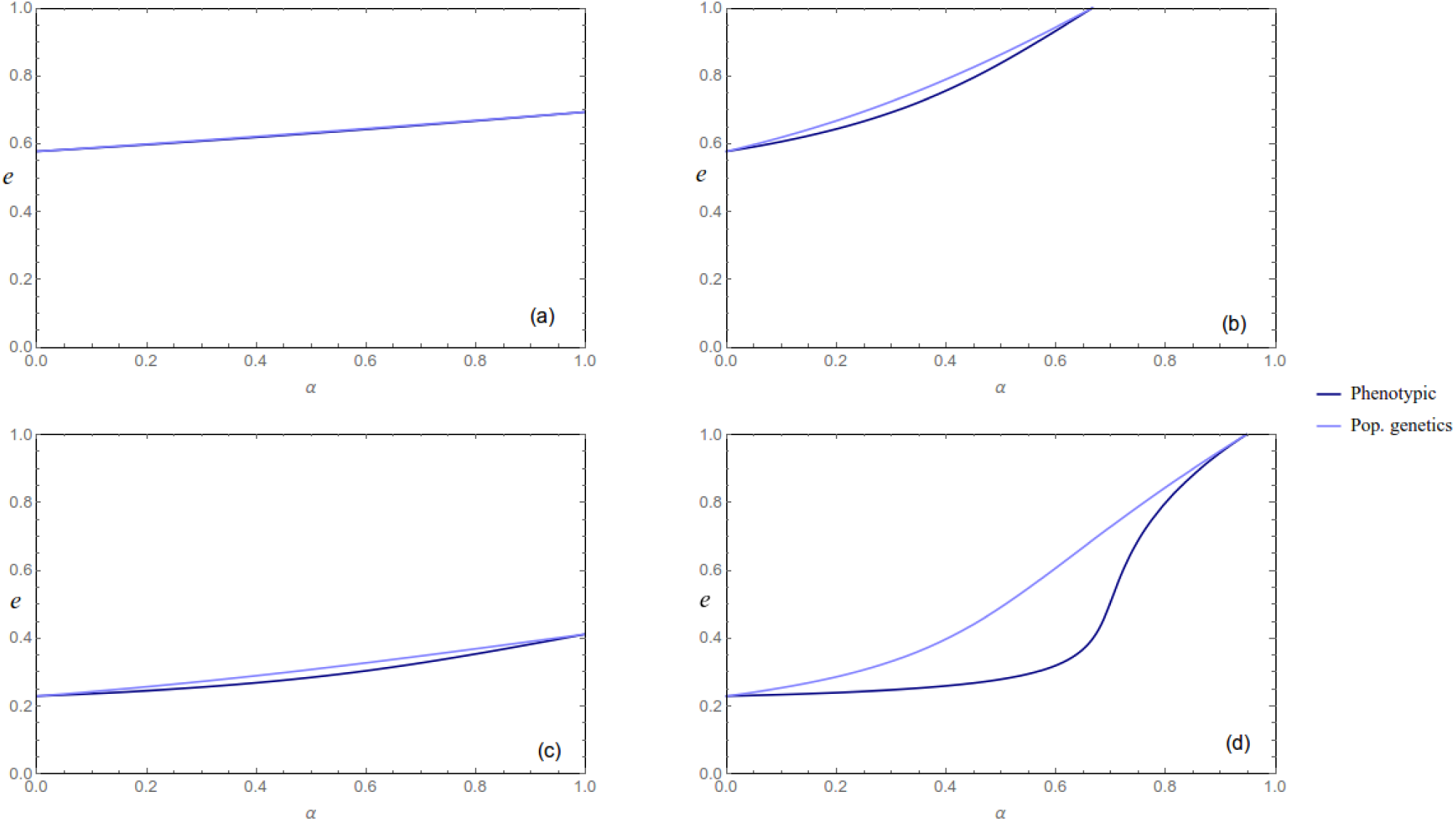
ESS reproductive effort is presented as a function of the selfing rate, in mild extrinsic mortality conditions (*S* = 0.75, Figures (a) and (b)), and in low extrinsic mortality conditions (*S* = 0.95, Fig. (c) and (d)), with low (*δ*_*a*_ = 0.1, Fig. and (c)) and high (*δ*_*a*_ = 0.1, Fig. (b) and (d)) adult inbreeding depression. The analytical expression we obtain with our main model is presented in light blue. The one we obtain with our phenotypic approach is presented in dark blue.

Using the same approach as for the evolution of lifespan, we now study the influence of lifespan for the evolution of self-fertilisation. For that matter, we do not need to specify any trade-off between survival and reproduction. Therefore, we will use the notation *𝒮*_*o*_(*e*) to denote survival probability, and fitness becomes

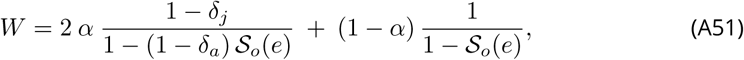

where the 2*α* term stands for Fisher’s transmission advantage [**fisher1941**]. The fitness of an individual taken at random in a fully outcrossing population (*W*_*o*_) is given by

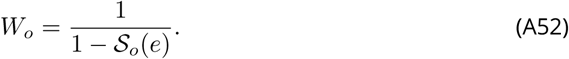

Such a population will be invaded by a self-fertilising mutant if

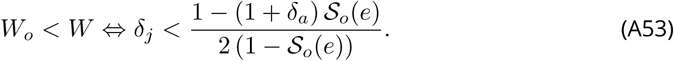

Furthermore, if this condition is satisfied, we have

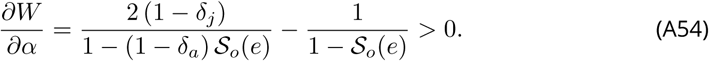

Therefore, only two evolutionarily stable strategies can exist, either *α* = 1 if Condition (A53) is satisfied, and *α* = 0 otherwise. These results completely match with those we obtained using our modifier model.

## APPENDIX V

### Numerical analyses

In this section, numerical analyses results are presented along with our analytical predictions.

**Figure A2.**
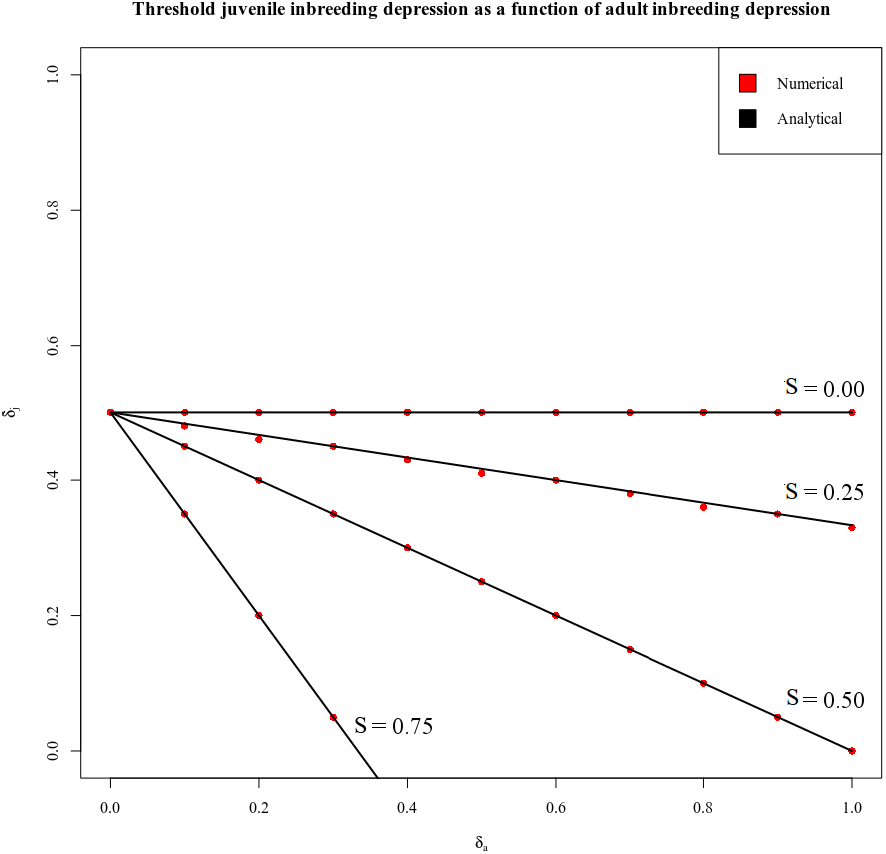
Numerical analyses results for the evolution of self-fertilisation. Threshold juvenile ID is presented as a function of adult ID, for various survival probabilities (*𝒮*_*o*_(*e*) = 0; 0.25; 0.5; 0.75).

For the evolution of self-fertilisation, the analysis was conducted starting from *α*_0_ = 0 and introducing mutants increasing the selfing rate. Good agreement was found between our analytical predictions and numerical analyses. Figure A2 shows some inbreeding depression threshold values obtained for various longevity situations. Either full selfing or strict outcrossing always evolved.

**Figure A3.**
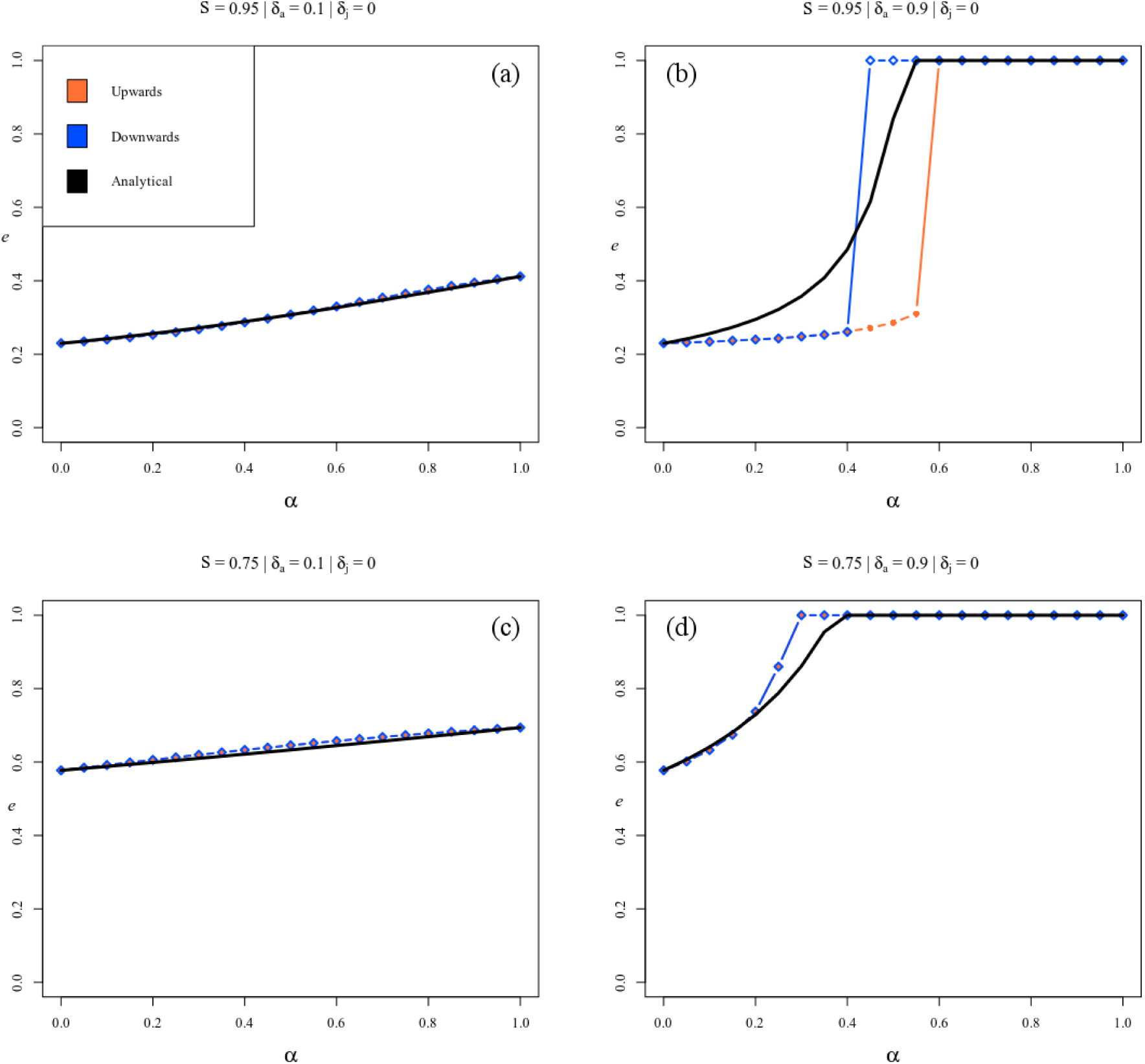
Numerical analyses results for the evolution of reproductive effort. ESS reproductive effort is presented as a function of the selfing rate, in low (*S* = 0.95, Fig. (a) and (b)) and mild (*S* = 0.75, Figures (c) and (d)) extrinsic mortality conditions, with low (*δ*_*a*_ = 0.1, Fig. (a) and (c)) and high (*δ*_*a*_ = 0.9, Fig. (b) and (d)) adult inbreeding depression. Numerical analyses are presented in orange (*upwards*) and blue (*downwards*). Analytical predictions are presented in black.

For the evolution of reproductive effort, two types of analyses were conducted: starting from *e*_0_ = 0 and introducing mutants increasing reproductive effort (*upwards* analysis, presented in orange), and starting from *e*0 = 1 and introducing mutants decreasing reproductive effort (*downwards* analysis, presented in blue). Part of the results are presented in Figure A3. Discrepancies between upwards and downwards analyses were observed in situations involving high adult inbreeding depression and low extrinsic mortality for intermediate selfing rates (Fig. A3b). Such situations are never reached in the context of the joint evolution of lifespan and selfing, because inbreeding depression is too high for selfing to evolve and intermediate selfing rates are never evolutionarily stable. Besides, good agreement was found between analytical predictions and numerical analyses in most cases, especially for selfing rates close to 0 or 1 (Fig. A3), which are the states that matter for the joint evolution of lifespan and selfing. Therefore, we considered our analytical prediction to satisfactorily agree with numerical analyses.

## APPENDIX VI

### Inbreeding depression affecting fecundity

In this appendix, we investigate the effect of inbreeding depression affecting fecundity, that is the contribution of selfed individuals to the gamete pool, on the coevolutionary dynamics. In Figure A4, we show the equilibria reached by simulations with different starting points, when setting *δ*_*j*_ = 0 and *δ*_*f*_ = 0.1; 0.4; 0.7 with *δ*_*a*_ = 0.4 and *S* = 0.8. Solid lines represent the analytical expectations in the absence of inbreeding depression on fecundity (*δ*_*f*_ = 0), and with *δ*_*j*_ = 0.1; 0.4; 0.7.

**Figure A4.**
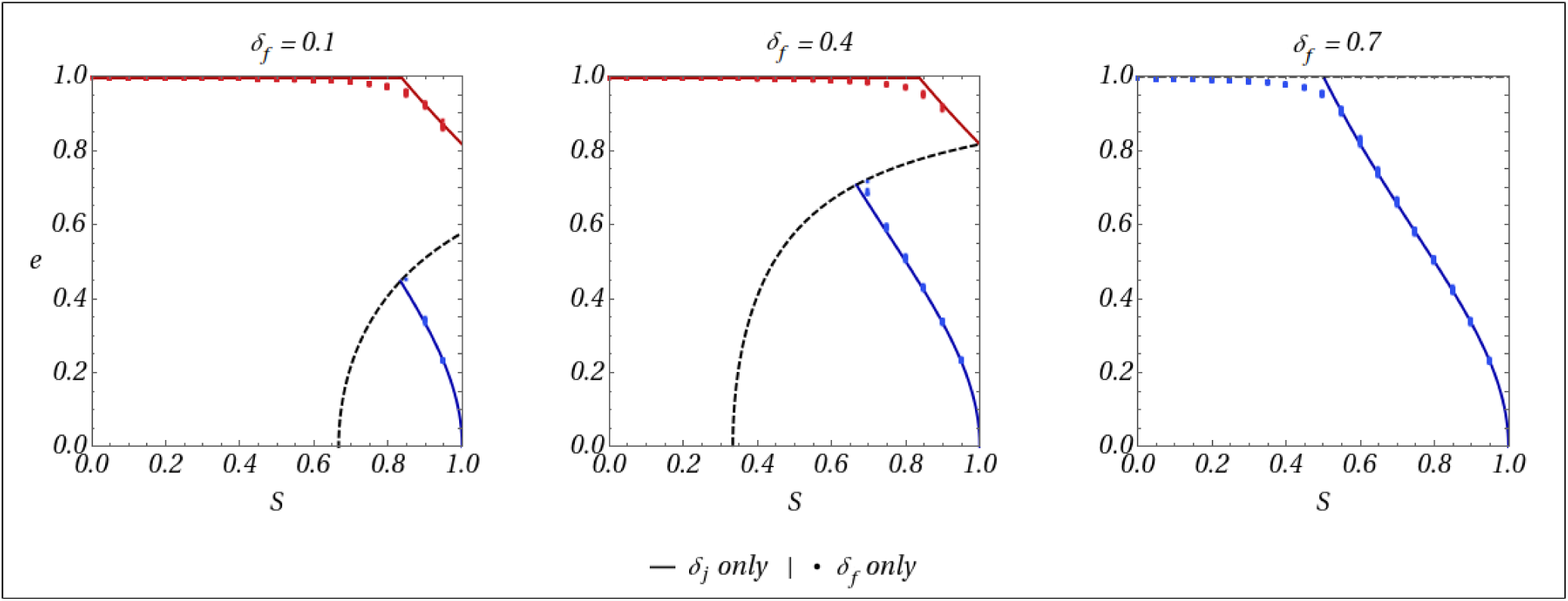
All results presented here assume *δ*_*a*_ = 0.1 and *x* = 2. Y-axis is the ESS reproductive effort (*e*^*\**^), and X-axis is extrinsic mortality (*S*). The red line corresponds to the equilibrium reproductive effort when *α* = 1 is reached. The blue line corresponds to the equilibrium reproductive effort when *α* = 0 is reached. The dashed black line represents the threshold lifetime inbreeding depression under which outcrossing can be maintained. Dots depict the results of individual-centered simulations. When red dots mean *α* = 1 is reached, while blue dots mean *α* = 0 is reached.

Simulations results are well predicted by these analytical expectations, which shows that increasing *δ*_*f*_ or *δ*_*j*_ has the same effect: it only reduces the range of conditions under which the full selfing strategy can evolve. The mechanisms underlying these similar behaviours are however slightly different. Indeed, while in both cases they are caused by a reduction of the participation of selfed individuals to reproduction, *δ*_*j*_ decreases the proportion of selfed individuals in the population. On the other hand, *δ*_*f*_ reduces the contribution of selfed adults to reproduction by reducing their contribution to the gamete pool.

